# In planta haploid induction in maize and tomato through disruption of KOKOPELLI

**DOI:** 10.64898/2026.06.16.731883

**Authors:** Nathanaël M.A. Jacquier, Jean-Philippe Mauxion, Andrea R.M. Calhau, Marina Millán Blánquez, Chloé Plagnard, Emilie Montes, Nathalie Gonzalez, Laurine M. Gilles, Thomas Widiez

## Abstract

Haploid induction is a key component of doubled haploid technology and an increasingly valuable tool for plant breeding, genome editing, and clonal seed production. While *in planta* haploid induction through haploid inducer lines offers significant advantages over *in vitro* approaches, its application remains limited in many crop species. Previously, disruption of the sperm cell-expressed *KOKOPELLI* (*KPL*) gene was shown to induce maternal haploids in *Arabidopsis thaliana*. Here, we report the creation of novel haploid inducer lines in two globally important crops, maize (*Zea mays*), a major staple food crop, and tomato (*Solanum lycopersicum*), a widely cultivated vegetable crop. Using targeted genome editing, we generated mutations in *KPL* orthologs and demonstrated that loss of *KPL* function confers haploid induction capacity, enabling the production of haploid seedlings in both species. These findings establish KPL as a conserved target to trigger haploid induction and expand the genetic toolbox available for haploid inducer development in crop species.

## Results

Haploid induction (HI) is an integral part of the double haploid technology and has traditionally been employed as a powerful tool in plant breeding to rapidly produce genetically fixed plant material (Jacquier *et al*., 2020). The utilization of HI has extended beyond its conventional role for doubled haploid production, now enabling the advancement of innovative methodologies such as haploid induction-editing to directly perform genome editing in commercial varieties (Kelliher *et al*., 2019) or as component of apomixis to produce clonal seeds (Underwood and Mercier, 2022). The induction of haploid embryos or seedlings entirely *in planta* through crosses with so-called haploid inducer lines represents an attractive strategy as compared to *in vitro* methods (Seguí-Simarro *et al*., 2021). However, this approach is not yet available in all crops and typically requires further optimization (Jacquier *et al*., 2020; Quiroz *et al*., 2024). Our previous work showed that disruption of the sperm cell expressed *KOKOPELLI (KPL)* gene from *Arabidopsis thaliana* (*AtKPL, AT5G63720*) is able to induce maternal haploid embryo in this non crop model plant (Jacquier *et al*., 2023). Here we performed gene editing of *KOKOPELLI* genes in both maize (*Zea mays*) and tomato (*Solanum lycopersicum*) and demonstrate that mutation of *KPL* is able to trigger HI in these two important crops.

Phylogenetic analyses identified KPL orthologs in different crop species and revealed KPL presence in both monocotyledonous and dicotyledonous (**Figure S1**) (Jacquier *et al*., 2023). Maize genome contains two paralogous *ZmKPL* genes: *Zm00001d002427/Zm00001eb071810, ZmKPLa* and *Zm00001d026242/ Zm00001eb430330, ZmKPLb*, whereas tomato genome contains a single *SlKPL* gene (Solyc05g008490) (**figure S1**). Gene expression data from publicly available transcriptomic dataset indicate specific flower expression of *SlKPL* and strong sperm cell enriched gene expression for *ZmKPLb* (**Figure S2-S3, Figure 1a**), consistent with Arabidopsis *AtKPL* expression found enriched in flower, pollen and sperm cell (Ron *et al*., 2010). *ZmKPLa* was found to be an extremely low expressed gene in the background noise level (inferior 0.2 FPKM in all tissue observed, **Figure S2**). Additional maize transcriptomic data from gamete and early embryo show specific sperm cell expression for both *ZmKPL*, with 142 times higher expression for *ZmKPLb* compared to *ZmKPLa* (**Figure 1a**). For maize, we used the transformable capacity of A188 inbreed line (WT) and our CRISPR-Cas9 editing tools (Doll *et al*., 2019) to mutate both *ZmKPLa* and *ZmKPLb* using a single construct. All selected CRISPR-Cas9 induced mutations led to a premature stop codon producing potentially truncated proteins of 28 amino-acids for *zmkpla-1* (503 amino acids in WT) and of 207- and 194 amino acids for *zmkplb-1* and *zmkplb-4* alleles respectively (706 amino acids in WT) (**Figure 1b**). To evaluate the HI capacity after inactivating KPL function in maize, pollen from each single mutant, as well as the double mutants *zmkpla-1 / zmkplb-4*, was used to pollinate a “glossy” female tester line carrying the *glossy1* mutation. This allows for easy identification of haploid seedling, which are scored by their ability to adhere water droplets due to the glossy phenotype (Gilles *et al*., 2017). Haploidy was further assessed by flow cytometry and haploid plants exhibited reduced size and sterility as compared to their diploid counterparts (**Figure 1c-1d**). Crosses with glossy female using *zmkpla-1* pollen led to 0.08% haploid induction rate (HIR), (1 haploid / 1269), whereas *zmkplb-1* and *zmkplb-4* single mutants led to 0.36% HIR (9/2472) and 0.73% HIR (4/551) respectively (**Figure 1e**). Crosses with glossy female using *zmkpla-1*/*zmkplb-4* double mutant pollen led to similar HIR as *zmkplb-4* single mutant (0.71% HIR, 23/3262) suggesting that loss of function of *ZmKPLb* is the main driver of haploid induction in the context of double mutant *zmkpla*/*zmkplb*. No haploid seedling was found when WT pollen were used (**Figure 1e**). For tomato, a genome editing construct harboring two SgRNA targeting the *SlKPL* gene was used to transform the WVa106 cultivar. *slkpl-1* CRISPR-Cas9 mutation was selected and induced a premature stop codon producing a potentially truncated protein of 22 amino-acids instead of 863 for the WT version (**Figure 1f**). *slkpl-1* mutant produced fruits with less seeds per fruit compared to the WT (**Figure 1g**). To evaluate the HI capacity of *SlKPL* inactivation in tomato, the DNA content of seedlings derived from self-pollinated homozygous *slkpl-1* mutant was assessed by flow cytometry and two haploids seedling were identified out of 761 screened (0.26% HIR), whereas no haploid seedling was found in self-pollinated WT plants (0/480) (**Figure 1h-i**). The haploid plants exhibited the typical reduced plant, leaf and flower size as compared to their diploid counterparts (**Figure 1j**).

**FIGURE 1.**
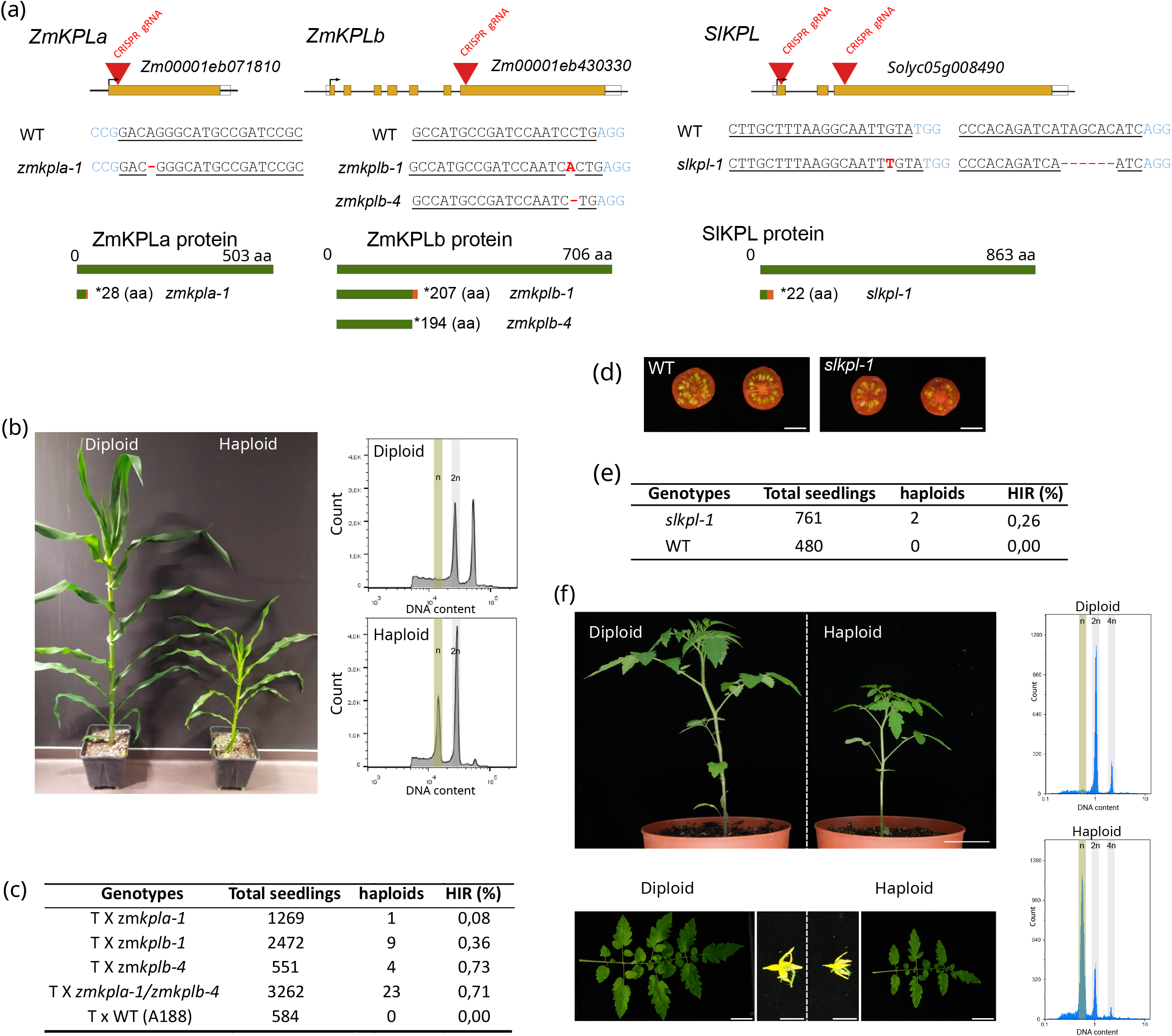
Loss of function of *KOKOPELLI* triggers haploid induction in maize and tomato. (a) Expression of *ZmKPLa* and *ZmKPLb* in maize gametes from (Chen et al., 2017). (b) *ZmKPLa* and *ZmKPL*b gene structures with CRISPR–Cas9 guide positions (red triangles). Exons (orange) and untranslated regions (empty square) are indicated. sgRNAs are underlined; PAM sequences are in blue and mutations in red. Wild-type protein sequences are shown in dark green, frameshift-derived sequences in dark orange. Asterisks indicate translation termination; peptide length is shown. (c) Two months old diploid vs haploid plants. (d) Representative illustrations of ploidy verification of maize haploid and diploid siblings using flow cytometry. (e) Haploid induction rate (HIR) of the different *zmkpl* lines tested. T: Glossy tester line. (f) *SlKPL* gene and protein structures with CRISPR–Cas9 guide positions, sgRNAs, mutations, and resulting protein effect. Legend as panel (b). (g) WT and *slkpl-1* tomato fruit. Scale bars: 1 cm (h) Illustrations of ploidy screening of tomato seedling using flow cytometry. (i) Haploid induction rate (HIR) of self-pollinated of *slkpl-1* mutant as compared to wild type (WT). (j) Tomato diploid and haploid plant phenotypes: 1-month-old whole seedling, 3-month-old leaves, and flowers. Bars: 5 cm for whole plants and for leaves and 1 cm for flowers.

In summary, targeted editing of *KOKOPELLI* in the monocot crop maize and the dicot vegetable crop tomato induces haploid embryo formation *in planta*. These findings, together with recent evidence that inactivation of *MtKPL* in *Medicago truncatula* also results in haploid induction (Wang *et al*., 2026), suggest that disruption of KPL function may represent a broadly applicable strategy for haploid induction across diverse plant species. However, haploid induction rates will need to be further improved for this approach to be compatible with breeding-scale deployment. Overall, our results identify KPL as a promising target for extending haploid induction technologies to a wider range of crops.

## Author Contributions

N.M.A. Jacquier, L. Gilles and T. Widiez conceived and designed the experiments; N.M.A. Jacquier, A.R.M. Calhau, M. Millán Blánquez, E. Montes and C plagnard conducted maize experiments and JP Mauxion conducted all tomato experiments; L. Gilles and N. Gonzalez contributed analytical tools. T. Widiez wrote the manuscript with contributions from all authors; L. Gilles and T. Widiez were involved in project management and PhD supervision of N.M.A Jacquier; T. Widiez obtained funding; T. Widiez and L. Gilles coordinated the project.

## Acknowledgements

We acknowledge the contribution of SFR Biosciences (UMS3444/CNRS, US8/Inserm, ENS de Lyon, UCBL) facilities, and in particular the help of E. Devevre with flow cytometry. We acknowledge the contribution of the Molecular Analysis Platform from Limagrain with genotyping and Priyanka Basavaraddi. The authors are thankful to Noah Vallois, Camille Knaupp, Justin Berger, Patrice Bolland, Isabelle Desbouchages, Hervé Leyral, Valérie Rouyère and Aurélie Honoré for technical assistance; Cindy Vial, Laureen Grangier, Nelly Camilleri, and Julie Prata for administrative assistance.

## Data Availability Statement

The data that supports the findings of this study are available in the Supporting Information of this article.

