## Supplemental material for "In planta haploid induction in maize and tomato through disruption of KOKOPELLI"

### Supplementary Methods

Plant materials and growth conditions

The maize (*Zea mays*) inbred line A188 and derived edited plants were grown under the French S2 safety standards for the culture of transgenic plants, in growth chambers or green house. The photoperiod consisted of 16 h light and 8 h darkness in a 24 h diurnal cycle. Temperature was set to 27°C/19°C (day/night). The relative humidity was controlled at 55% (day) and 65% (night). For tomato, (S. *lycopersicum*) cultivar WVa106 and derived edited *Slkpl* CRISPR-Cas9 mutants were grown in soil under long-day photoperiods (15:9). Daytime and night-time temperatures were 25–29°C and 17–19°C, respectively.

Maize transformation and gene editing

A. tumefaciens strain LBA4404 harboring the construct of interest was used to transform 12 DAP immature embryos of maize inbred line A188 according to a standard protocol (Fierlej *et al.*, 2022; Ishida *et al.*, 1996). Single guide RNAs (sgRNA) were designed using the CRISPOR web tool (Concordet and Haeussler, 2018) and two 20-nucleotide targets sequence were selected for each gene (**Supplemental Table S1**). Genome editing-related cloning steps were performed following the strategy described in (Doll *et al.*, 2019). Screening for genome editing events was performed on gDNA from T0 leaves seedling by specific PCR amplification (**Supplemental Table S1** for primer sequences) followed by Sanger sequencing.

Tomato transformation and gene editing

The CRISPR–Cas9 construction, plant transformation and obtention of homozygous mutated lines were performed as described in (Beauchet *et al.*, 2024). The same strategies as for maize were used for sgRNA design leading to sgRNA listed in (**Supplemental Table S1**). Screening for genome editing events of *Slkpl* CRISPR-Cas9 mutants was performed on gDNA from T0 leaves seedling by specific PCR amplification (**Supplemental Table S1** for primer sequences) followed by Sanger sequencing.

Pre-scoring of maize haploid induction

The maize *zmkpl* mutant lines to be tested for haploid induction were crossed as male parent with "glossy tester line" homozygous for the *glossy1* mutation (Gilles *et al.*, 2017; Melchinger *et al.*, 2016). Offspring's kernels was germinated and scored for the glossy (bright leaf surface, adhering water droplets) phenotype indicative of haploid plantlets. Haploid induction rate was determined as the percentage of glossy among germinated plantlets. Haploid state of glossy plantlets was double-checked by flow cytometry.

Flow cytometry analysis

For maize, youngest leaves from 2 weeks old plants were harvested (2cm2 in total). Leaves were finely chopped on glass Petri dishes and in 500µl of fresh and cold Galbraith's buffer (Galbraith et al., 1983). Glass dish were inclined to separate chopped leaves from the extract and additional 500µl of fresh and cold Galbraith's buffer were added on chopped leaves to drag additional nuclei down. Extract were filtered through 70µm, 40µm and 20µm nylon sieves on ice. Finally, 500ul of filtered extract is transferred in polystyrene (Falcon® 5 mL Round-Bottom Polystyrene Tube, ref: 352052) tube and 1.25µl of DAPI (5mg/ml) were added. Samples were directly analyzed in LSR Fortessa flow cytometer operating at 640nm (Red laser) and 405nm (violet laser) using respectively APC (670/14) and DAPI (450/50) filters. Cleaning and rinsing steps were performed between each sample. For tomato, young leaves from 3 weeks old plants were harvest (1cm$^2$ in total) in 96 well plate and flash freeze for storage. The plant tissues were slightly grinded in 200 µl of chilled CyStain UV Precise P Nuclei Extraction Buffer (Sysmex) using a Retsch grinder. Extracts were stained by adding 800 µl of chilled CyStain UV Precise P Staining Buffer (Sysmex) and the suspension were filtered through a 50 µm nylon filter. The nuclei DNA content was measured using a CyFlow Space flow cytometer (Sysmex) operating at 365nm (violet laser) using band pass filter (455/25). Cleaning and rinsing steps were performed between each sample.

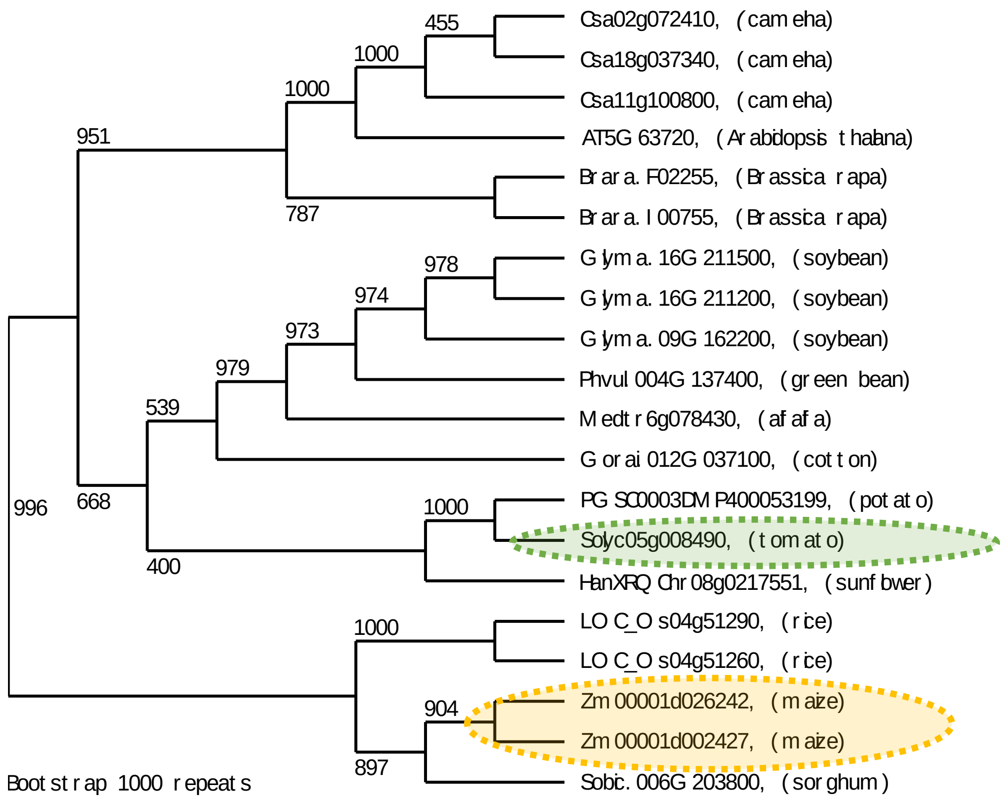


**<u>Figure S1</u>** : Phylogenetic tree of KOKOPELLI proteins in selected crops highlighting the two maize ZmKPL proteins (orange) and the tomato SlKPL protein (green), adapted from Jacquier *et al.*, 2023, 193, 1, 182–185.

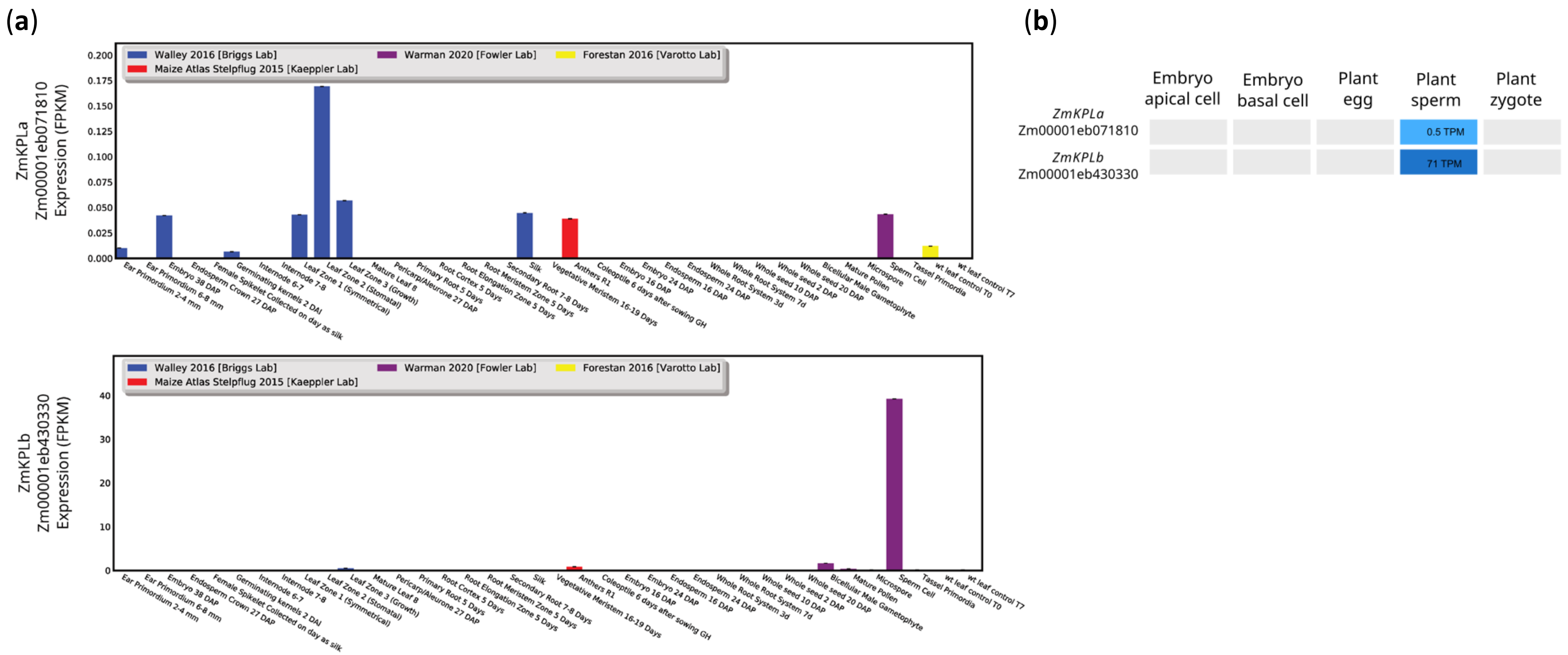


**Figure S2** : *ZmKPLa* and *ZmKPLb* gene expression data from from publicly available transcriptomic dataset.

(a) Data are from qTeller resource Woodhouse et al., 2022, *Bioinformatics*, 38, 1, 236–242.

(b) Expression of *ZmKPLa* and *ZmKPLb* in maize gametes from Chen et al., 2017, Plant cell, 29, 9, 2106–2125.

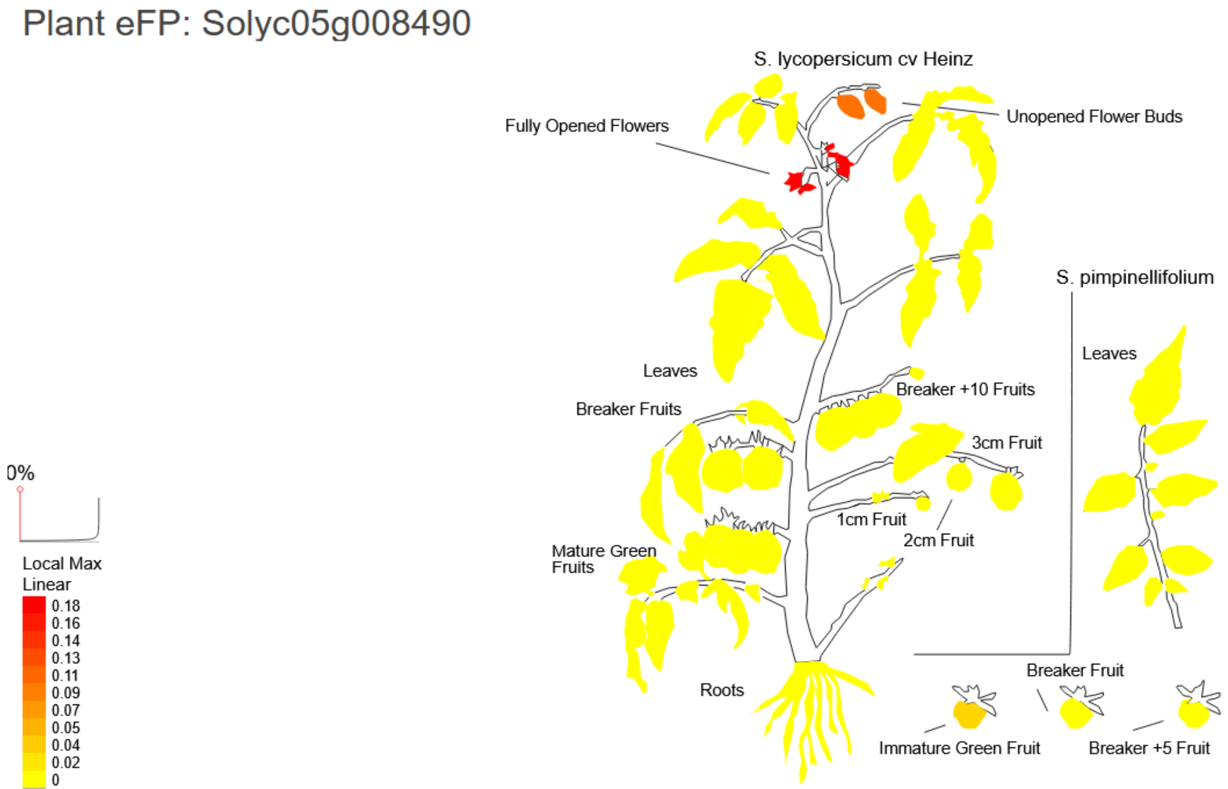


**Figure S3** : *SlKPL* gene expression data from transcriptomic analyses.
This image was generated with the Plant eFP at bar.utoronto.ca/eplant by Waese et al. 2017. Data are from The tomato genome sequence providing insights into fresh fruit evolution: Tomato Genome Consortium, 2012, *Nature* **485**, 635–641.

**Supplemental Table S1**: sgRNA sequences (5′–3′; PAM underlined) and primer sequences (5′–3′) used for th

| | sequences |
|---|---|
| sgRNA_1_zmkpla | CCGGACAGGGCATGCCGATCCGC |
| sgRNA_2_zmkpla | CTTAACCTGCAACACTCTACGG |
| sgRNA_1_zmkplb | CCATCAGGTTCTCTAATGTCCCC |
| sgRNA_2_zmkplb | GCCATGCCGATCCAATCCTGAGG |
| sgRNA_1_Slkpl1 | CTTGCTTTAAGGCAATTGTATGG |
| sgRNA_2_Slkpl1 | CCCACAGATCATAGCACATCAGG |
| ZmKPLa_Mu1_2_F1 | GTAATGCAAACAGTCGCTTTGTTC |
| ZmKpla_Crispr_R1 | GTTCTCGACGAGCGATTGAAG |
| ZmKPLb_F1 | CAGGGCTCTTCTGATGAAGATACTA |
| ZmKPLb_CRISPR_R1 | GTTCCAGCTACGGGGAAACG |
| SlKPL-F1 | GGTAGTTGTAGCAGTGTAGTTG |
| SlKPL-R1 | GGTAATGGTGGAAGTTACGAG |
| SlKPL-F2 | GGACCATGAAGCGCGAGTCAT |
| SlKPL-R2 | CGATCTCCATCGCATATCAAG |

ıe generation and genotyping of edited plants, respectively.
